# UniProt Genomic Mapping for Deciphering Functional Effects of Missense Variants

**DOI:** 10.1101/192914

**Authors:** Peter B. McGarvey, Andrew Nightingale, Jie Luo, Hongzhan Huang, Maria J. Martin, Cathy Wu, the UniProt Consortium

**Author notes:** Email addresses: PBM^§^ AN, J.L, MJM, H.H, CW. Grant Sponsor: NIH/NIGRI grants U41HG007822, U41HG007822-02S1 for UniProt and U01HG007437 for ClinGen.

## Abstract

Understanding the association of genetic variation with its functional consequences in proteins is essential for the interpretation of genomic data and identifying causal variants in diseases. Integration of protein function knowledge with genome annotation can assist in rapidly comprehending genetic variation within complex biological processes. Here, we describe mapping UniProtKB human sequences and positional annotations such as active sites, binding sites, and variants to the human genome (GRCh38) and the release of a public genome track hub for genome browsers. To demonstrate the power of combining protein annotations with genome annotations for functional interpretation of variants, we present specific biological examples in disease-related genes and proteins. Computational comparisons of UniProtKB annotations and protein variants with ClinVar clinically annotated SNP data show that 32% of UniProtKB variants co-locate with 8% of ClinVar SNPs. The majority of co-located UniProtKB disease-associated variants (86%) map to ‘pathogenic’ ClinVar SNPs. UniProt and ClinVar are collaborating to provide a unified clinical variant annotation for genomic, protein and clinical researchers. The genome track hubs, and related UniProtKB files, are downloadable from the UniProt FTP site and discoverable as public track hubs at the UCSC and Ensembl genome browsers.

## Introduction

Genomic variants may cause deleterious effects through many mechanisms, including aberrant gene transcription and splicing, disruption of protein translation, and altered protein structure and function. Understanding the potential effects of non-synonymous (missense) single nucleotide polymorphisms (SNPs) on protein function is a key for clinical interpretation (MacArthur et al., 2014; Richards et al., 2015). UniProtKB, which represents decades of effort in literature-based and semi-automated expert protein curation, contains a wealth of information that is potentially valuable for missense variant annotation, including positional information on enzyme active sites, modified residues, and binding domains, as well as phenotypic consequences of sequence variants (M. L. Famiglietti et al., 2014). Currently for the 20,396 human reviewed entries in the UniProtKB/Swiss-Prot section, 99.9% have at least one sequence feature including a structural or functional domain, 63% have variant annotation, 31% have one or more 3-D structure, and 24% have an annotated post translational modification. Aligning curated protein functional information with genomic annotation and making it seamlessly available to the genomic, proteomic and clinical communities should greatly inform studies on the functional consequences of variants. Although UniProtKB is being exploited for this purpose to small extent, for example PolyPhen-2 (Adzhubei, Jordan, & Sunyaev, 2013) incorporates information on protein active sites from UniProtKB, it is vastly underutilized. PolyPhen and other tools use a variety of structural and sequence conservation information to predict the effects of missense variants and have been incorporated into variant interpretation resources and commercial pipelines (Ioannidis et al., 2016; Kircher et al., 2014; McLaren et al., 2016; Nykamp et al., 2017; Qian et al., 2018; Shihab et al., 2013; Shihab et al., 2015). While these tools work very well for some well-studied genes (e.g., *BRCA1, TP53)*, the results are less established for many others and improvements are needed (Guidugli et al., 2018; Karbassi et al., 2016; Mahmood et al., 2017; Qian et al., 2018; Tavtigian et al., 2018).

Genome browsers (Kent et al., 2002; Yates et al., 2016) provide an interactive graphical representation of genomic data. They utilize standard data file formats, enabling the import and integration of multiple independent studies, as well as an individual user’s own data, through community track hubs (Raney et al., 2014). Here, we illustrate the utility of representing UniProtKB protein functional annotations at the genomic level via track hubs and demonstrate how this information can be used in combination with genomic annotations to interpret the effect of missense variants in disease-related genes and proteins using specific biological examples and some larger scale comparisons.

Knowledge of a variant’s disease associations is also important in evaluating its impact. Many resources, including UniProtKB and ClinVar (Landrum et al., 2018) provide disease-related information on variants. In UniProtKB, the majority of this information comes from the literature, although OMIM (OMIM, 2018) is also used, primarily as a source for disease names and descriptions and as a means identifying relevant literature. ClinVar is an open database for the deposition of variants identified in clinical genome screens; the scientist submitting variant information is responsible for assigning a clinical significance class to individual variants following the ACMG clinical significance recommendations (Richards et al., 2015). A subset of ClinVar’s variations, non-synonymous SNPs that change a single amino acid, closely reflects UniProtKB’s “Natural variants”, which include polymorphisms, variations between strains, isolates or cultivars and disease-associated mutations (https://www.uniprot.org/help/variant) and are mostly (98%) single amino acid changes. We evaluated UniProt Natural variant annotation against equivalent annotations in co-located ClinVar SNPs and found significant synergy between the two resources.

## Methods

Mapping UniProtKB protein sequences to their genes and genomic coordinates is achieved with a four phase Ensembl import and mapping pipeline. The mapping is currently conducted for the UniProt human reference proteome with the GRCh38 reference sequence and also for *Saccharomyces cerevisiae* S288C with the sacCer3 reference sequence. Reference sequences are provided by Ensembl. We summarize the approach here. Additional details, figures and references are provided in the Supplemental Methods and Results document.

### Phase One: Mapping Ensembl identifiers and translations to UniProtKB sequences

UniProtKB imports Ensembl translated sequences and associated identifiers, including gene symbols and the HGNC identifier. An Ensembl translation is mapped to a UniProtKB sequence only if the Ensembl translated sequence is 100% identical to the UniProtKB sequence with no insertions or deletions. When an Ensembl translation does not match an existing UniProtKB canonical sequence or an isoform in a UniProtKB/Swiss-Prot entry, the Ensembl translation is added as a new UniProtKB/TrEMBL entry.

### Phase Two: Calculation of UniProt genomic coordinates

Given the UniProt to Ensembl mapping, UniProt imports the genomic coordinates of every gene and the exons within a gene. Included are the 3’ and 5’ UTR offsets in the translation and exon splice phasing. With this collated coordinate data, UniProt calculates the portion of the protein sequence in each exon and defines the genomic coordinates for the amino acids at the beginning and end of each exon. This set of peptide fragments with exon identifiers and coordinates is stored as the basis for protein to genomic mappings in UniProt.

### Phase Three: Converting UniProt position annotations to their genomic coordinates

UniProtKB sequence position annotations or “features” have either a single amino acid location or amino acid range within the UniProtKB canonical protein sequence. Using the exon coordinates of the protein peptide fragments, the genomic coordinates of a feature annotation are calculated by finding the amide (N) terminal exon and the carboxyl (C) terminal exon. A range of all positions from the first nucleotide in the first amino acid codon through to the last nucleotide position in the last amino acid codon are mapped. Details and figure in Supplemental Methods.

### Phase Four: UniProt BED and BigBed Files

Converting protein functional information into its genomic equivalent requires standardized formats. The Browser Extensible Data (BED) (UCSC, 2016a), a tab-delimited format, represents one format for displaying UniProtKB protein annotations on a genome browser. The binary equivalent of the BED file is BigBed (Kent, Zweig, Barber, Hinrichs, & Karolchik, 2010); this format is more flexible in allowing additional data elements, providing a greater opportunity to fully represent protein annotations and is one of the file formats used to make track hubs. A track hub is a web-accessible directory of files that can be displayed in track hub-enabled genome browsers (Raney et al., 2014). Hubs are useful, as users only need the hub URL to load all the data into the genome browser.

Moreover, a public registry for track hubs is now available (https://trackhubregistry.org/) allowing users to search for track hubs in one location and providing links to multiple genome browsers.

Using the protein genomic coordinates, with additional protein feature specific annotations from UniProtKB, the BED detail (UCSC, 2016b) and BigBed formatted files, as well as track hub required files, are produced for the UniProtKB human reference proteome. Genomic coordinates are converted to the zero-based coordinates used within the BED file formats

### Mapping ClinVar SNPs to protein features and variants

Data for comparing ClinVar SNPs to UniProtKB features comes from the ClinVar variant_summary file from NCBI (ftp.ncbi.nlm.nih.gov/pub/clinvar/tab_delimited/variant summary.txt.gz), the UniProtKB feature specific BED files (ftp.uniprot.org/pub/databases/uniprot/currentrelease/knowledgebase/genome annotation trac ks/UP000005640 9606 beds/) and the human variation file on the UniProt FTP site: (ftp.uniprot.org/pub/databases/uniprot/current_release/knowledgebase/variants/humsavar.txt).1) For each feature in UniProtKB, we check the genomic position against the position for each SNP in ClinVar. If the genome positions of the protein feature overlap the chromosome and genomic coordinate of the SNP we establish a mapping. Information about the SNP and the feature, including the amino acid change are attached to the mapping file. 2) For each result in 1, we check the SNP position against the exon boundary for the protein. A flag is added if a SNP coordinate is within the exon boundary. Variants outside of exons were excluded from further analysis here. 3) For historical reasons, disulfide bonds are annotated in UniProtKB as a range between the two cysteines that form the bond. For comparison with ClinVar, we extract the positions of the first and last cysteines from this range as only variants at these two positions are expected to affect bond formation. 4) For each UniProt variant in 2 and 3, we check that the ClinVar provided RefSeq accession and UniProt accession refer to the same sequence and check that the amino acid change reported in UniProt and ClinVar is the same. 5) To compare Pubmed IDs (PMIDs) cited as evidence by the two resources, we use the following files: (i) for ClinVar: ftp.ncbi.nlm.nih.gov/pub/clinvar/tab_delimited/var_citations.txt from NCBI and (ii) for UniProtKB: ftp.uniprot.org/pub/databases/uniprot/current_release/knowledgebase/complete/uniprot sprot.d at.gz. For each co-located variant, we use the ClinVar Allele ID and UniProtKB Variant ID to extract the relevant PMIDs for each variant and construct a table of all the PMIDs and PMID counts.

### Comparison of UniProt and ClinVar Variant Annotation

UniProt classifies variants into three categories: 1) Disease – variants reported to be implicated in disease; 2) Polymorphism – variants not reported to be implicated in disease; 3) Unclassified – variants with uncertain implication in disease as evidence for or against a pathogenic role is limited, or reports are conflicting. ClinVar does not annotate variants directly but accepts assertions made by submitters with documentation on their criteria and associated documents and publications (if any) and classifies them into groups based on levels of evidence (0-4 gold stars). The predominant assertions in ClinVar, which are the ones we used for comparison, are those recommended by the ACMG/AMP guidelines (Richards et al., 2015): Benign, Likely benign, Uncertain significance, Likely pathogenic and Pathogenic. There are a small number of additional disease related assertions in ClinVar such as ‘risk factor’ and ‘drug response’, which we classified as ‘other’ in our analysis. All of the ClinVar assertions in the ‘other’ category that aligned with UniProt annotations were ‘drug response’ assertions. We only used variants with 1-4 stars and removed all 1-star variants with conflicting interpretations and those with no associated phenotype. We equated ClinVar assertions to UniProt classifications as follows: all ‘pathogenic’ assertions (Pathogenic and Likely pathogenic) to ‘Disease’ in UniProt; ‘Uncertain significance’ to ‘Unclassified’; and, all ‘benign’ (Benign and Likely benign) assertions to ‘Polymorphism’.

## Results

### Mapping UniProtKB human protein annotations to the Genome Reference (GRC)

Functional positional annotations (called Features) from the UniProt human reference proteome are now being mapped to the corresponding genomic coordinates on the GRCh38 version of the human genome for each release of UniProt. These mappings are available as BED files or as part of a UniProt genomic track hub and can be downloaded from the UniProt FTP site (www.uniprot.org/downloads). In addition, they are discoverable as a public track hub in the UCSC and Ensembl genome browsers by searching for UniProt. The track hub has been registered with the track hubs registry (trackhubregistry.org/search?q=uniprot&type=genomics), which provides links to load the tracks in either browser. For the UniProt 2018_01 release, the locations of 112,093 human reference protein sequences from UniProtKB were mapped to the GRCh38. This includes 18,687 canonical and 14,783 isoform sequences from UniProtKB/Swiss-Prot and 56,363 sequences from UniProtKB/TrEMBL. Thirty-four different positional annotation types (e.g. active sites, modified residues, domains and amino acid variants) with associated information curated from the literature are currently aligned with the genome sequence. **Table 1** shows the full list of positional annotations (features) and the number of each feature currently mapped to the human genome.

**Table 1:**
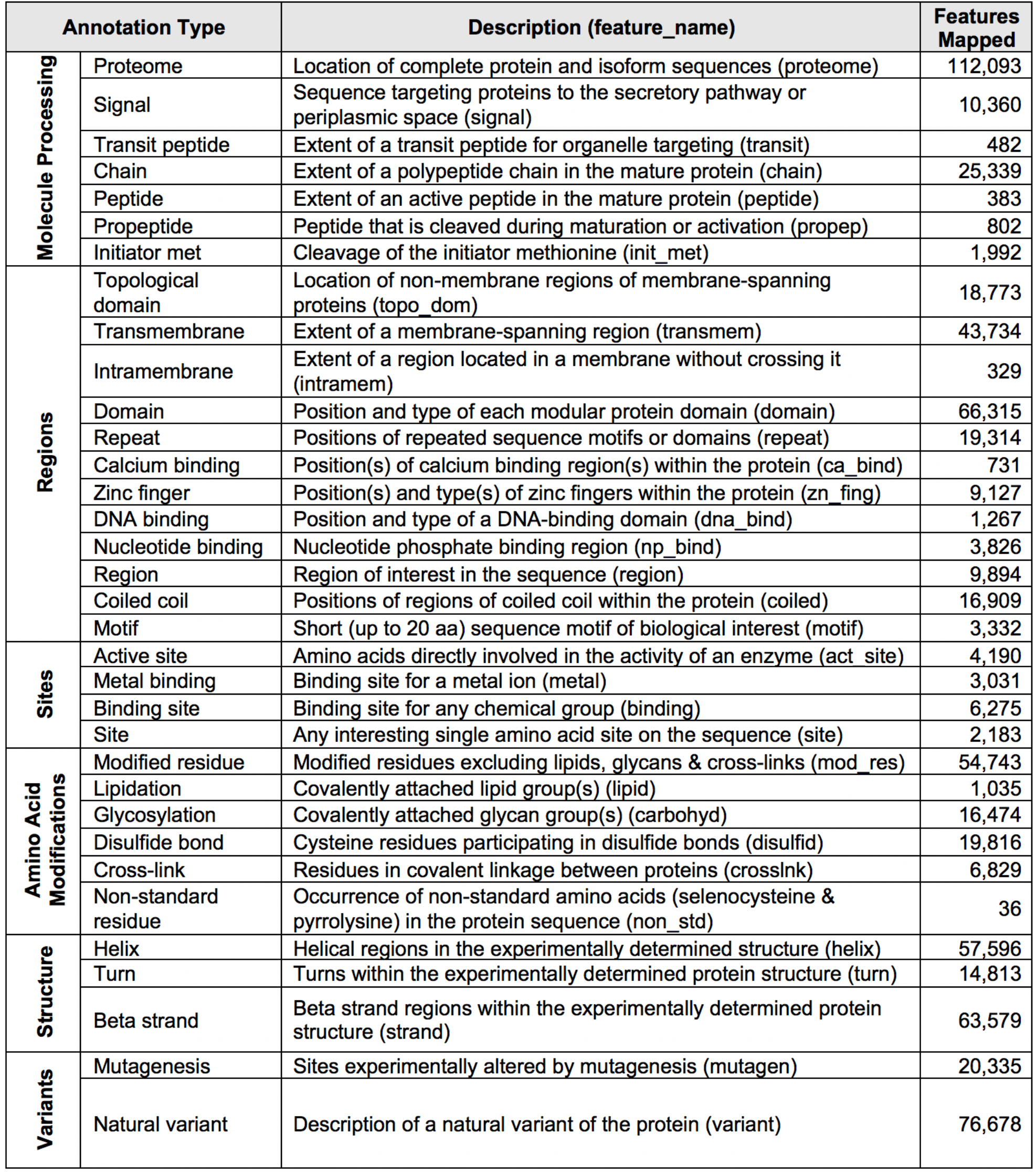
UniProtKB sequence annotations in track hubs. Annotation types, descriptions and current number of each feature mapped to the human genome are shown. UniProtKB release 2018_01 (Jan 2018) was used for this table. For more information on sequence features in UniProtKB see (www.uniprot.org/help/sequence_annotation).

### Coverage

All Ensembl human proteome translations are mapped to locations on the genome. However, not all UniProt human proteins are mapped. Because we require 100% identity between the Ensembl and UniProt sequences, a relatively small number of well annotated proteins in UniProtKB/Swiss-Prot are not mapped to the current genome (∼5%). Some of these proteins are uncharacterized proteins, endogenous retrovirus proteins, and non-germline sequences related to cancer or the immune system, which could be considered a lesser priority.

However, others are experimentally characterized proteins that use sequences that do not match the current consensus build. In some cases, the differences are small and can be addressed by curation or relaxing the match criteria. Other proteins such as mucin proteins encoded by the genes *MUC2* and *MUC19* are known to have variable repeat regions and the variation between the experimentally curated sequence and the current reference sequence is much greater. We are developing processes to map some of these and will make them available in future releases. In addition, some proteins map to multiple locations. Duplicate mappings occur because: 1) Gene duplications produce identical protein sequences. As the functional and structural literature predates the human genome, UniProtKB has chosen to maintain only one entry for some proteins mapped to multiple genes (for example, see UniProtKB P69905, hemoglobin subunit alpha protein encoded by the *HBA1* and *HBA2* genes, and UniProtKB Q16637, the survival motor neuron protein encoded by *SMN1* and *SMN2).* 2) Some genes/proteins map to chromosomal regions with multiple alternative assemblies which are included in our data. For example, UniProtKB P28068, HLA class II histocompatibility antigen, DM beta chain encoded by the *HLA-DMB* gene has one primary and seven alternate mappings (https://www.ensembl.org/Human/Search/Results?q=P28068). 3) Homologous genes are found on pseudoautosomal regions of the X and Y chromosome (Helena Mangs & Morris, 2007; Veerappa, Padakannaya, & Ramachandra, 2013).

### Usage

The BED tab-delimited files are useful to extract genome locations and annotation for data integration and computational analysis similar to that described below for mapping to ClinVar SNPs. However, we recommend using track hubs and not the BED text files on genome browsers. The extended BED format is not supported in a consistent manner on all browsers but the track hubs (BigBed format) are supported and provide enhanced functionality. The track hubs are set by default to provide ten feature tracks. Not all track hub enabled browsers are correctly interpreting this option; depending on which browser you use, you may need to turn on or off the feature tracks you prefer in in the browser controls.

### Biological Examples

To illustrate the utility of combining UniProt protein feature annotations and variation annotations to determine a probable mechanism of action, we looked at two well-studied disease-associated proteins. The alpha-galactosidase A *(GLA)* gene (HGNC:4296, UniProtKB P06280) has been linked to Fabry disease (FD) (MIM# 301500) (Romeo & Migeon, 1970; Schiffmann, 2009), a rare X-linked lysosomal storage disease where glycosphingolipid catabolism fails and glycolipids accumulate in many tissues from birth. Many of the protein-altering variants in *GLA* are associated with FD. UniProt curators have recorded 220 amino acid variants and 6 deletions, of which, 219 are associated with FD. ClinVar has 193 SNPs and 34 deletions in *GLA* associated with FD, of which 155 have an assertion of Pathogenic or Likely pathogenic. One hundred and four of the ClinVar SNPs align with 100 UniProt amino acid variants and cause the same amino acid change. These variants are distributed evenly over the entire protein sequence. **Figure 1** shows a portion of exon 5 of the *GLA* gene on the UCSC genome browser. <u>**Panel 1 selection A**</u> illustrates a situation where an amino acid that is part of the *GLA* active site is affected by a variant (P06280:p.Asp231Asn (<u>uniprot.org/uniprot/P06280#VAR_000468</u>)). The acidic proton donor aspartic acid (Asp, D) is replaced by the neutral asparagine (Asn, N) suggesting it no longer functions properly as a proton donor in the active site. This missense variant is associated with Fabry disease (Redonnet-Vernhet et al., 1996). **<u>Selection B</u>** shows a cysteine (Cys, C) residue annotated by UniProt as participating in a disulfide bond that aligns with a FD-associated missense variant (P06280:p.Cys223Gly (<u>uniprot.org/uniprot/P06280_#VAR_012401</u>); (Germain & Poenaru, 1999)). The variant converts a cysteine to a glycine, resulting in the loss of the wild-type disulfide bond between a β-strand and the C-terminal end of an a-helix encoded by exons four and five. Disruption of the disulfide bond disrupts the structure of the protein and is an obvious mechanism of action for the pathogenicity of this variant. Currently the P06280:p.Cys223Gly variant is not annotated in ClinVar or dbSNP. Note, seven of the ten cysteines involved in the five disulfide bonds in alpha-galactosidase A have annotated variants associated with the disease. A comparison of all ClinVar SNPs that overlap cysteines that are annotated as forming disulfide bonds in UniProtKB showed that 86% were annotated as pathogenic, a higher proportion than for any other feature in UniProtKB. See figure 2 and discussion below. **<u>Figure 1 Panel 2 selection C</u>** shows an N-linked glycosylation site overlapping multiple missense variants annotated in ClinVar as pathogenic (www.ncbi.nlm.nih.gov/clinvar/variation/10730/) and in UniProtKB as associated with FD (P06280:p.N215S (<u>uniprot.org/uniprot/P06280#VAR_000464</u>)). The P06280:p.Asn215Ser variant has been described in the literature multiple times. In ClinVar, multiple submitters have annotated this variant as pathogenic, citing twenty-four publications describing patients and family pedigrees with FD. In UniProtKB, the variant has five associated publications, three also found in ClinVar and two unique. In addition, UniProtKB cites two publications (<u>uniprot.org/uniprot/P06280#ptm_processing</u>) that review the 3-D structure and molecular details of the function of glycosylation at this and other sites (Chen et al., 2009; Garman & Garboczi, 2004). Evidence associated with the UniProtKB “Glycosylation” feature shows that the oligomannose-containing carbohydrate at this Asn215 site plus the Asn192 site (not shown) are responsible for secretion of the active enzyme (Ioannou, Zeidner, Grace, & Desnick, 1998) and targeting to the lysosome (Ghosh, Dahms, & Kornfeld, 2003). Mutation of Asn215 to Ser eliminates the carbohydrate attachment site, causing inefficient trafficking of the enzyme to the lysosome.

**Figure 1.**
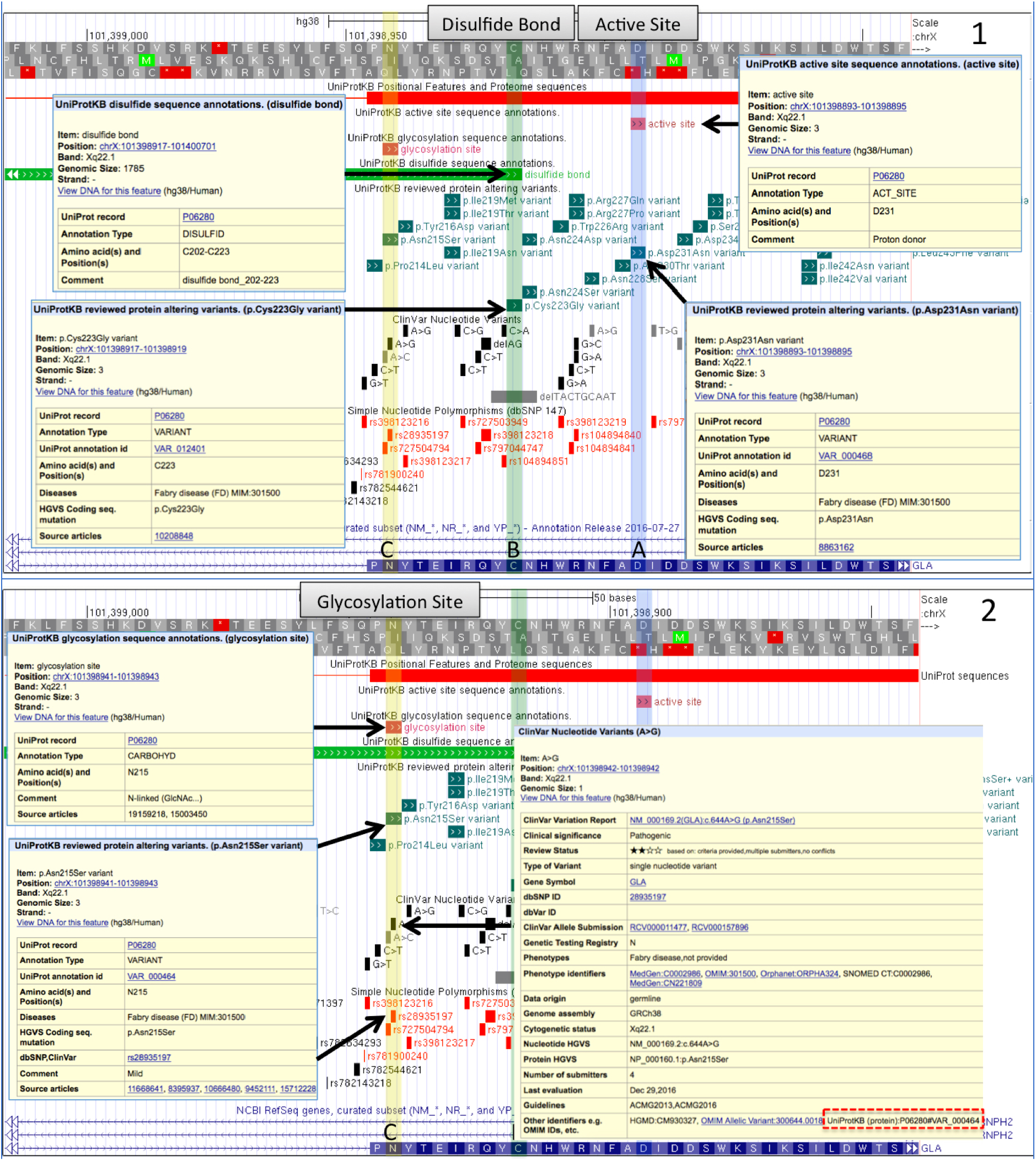
The *GLA* gene (P06280, alpha-galactosidase A) associated with Fabry disease (FD) shown on the UCSC browser with UniProtKB genome tracks plus ClinVar and dbSNP tracks. **Panel 1 selection A** shows UniProt annotation for part of the enzyme’s Active Site and an amino acid variation from a SNP associated with FD that removes an acidic proton donor (Asp, D) is replaced by the neutral (Asn, N). In selection B another variant disrupts an annotated disulfide bond by removing a cysteine required for a structural fold. SNPs are not observed in the other data resources in these positions. **Panel 2 selection C** shows an N-linked glycosylation site disrupted by another UniProt amino acid variant that does overlap pathogenic variants in ClinVar and other databases. Links between UniProt and ClinVar are illustrated in the display.

A second biological example where variants associated with Alzheimer disease disrupt enzymatic cleavage sites for peptides found in toxic amyloid plaques in the brains of Alzheimer patients is described in the supplemental methods and results.

In these examples, we looked at the annotation of individual variants manually but, as we illustrate below, our alignment of genome and protein variant annotations can be applied to larger scale analyses.

### UniProt Features containing ClinVar SNPs

As illustrated above, a missense variant in a key functional feature of a protein may alter a protein’s structure and function and if severe enough might be classified as harmful. To get an overview of variants in different functional features we examined SNPs from ClinVar that overlap selected protein features. For this comparison, we grouped the five category ACMG/AMP assertions into three 1) pathogenic, 2) uncertain significance, and 3) benign and only considered ClinVar SNPs with 2-4 gold stars and selected SNPs with one gold star (see Methods). **Figure 2** plots the percentage of ClinVar SNPs in each annotation category that overlap in each feature type (Original data in Supplemental Methods). Six features have more pathogenic variant classifications than either benign or uncertain (Disulfide Bonds, Initiator Methionine, Intramembrane Region, Natural Variant, DNA Binding Domain, Active Site). For three features (Nucleotide binding region, Lipid attachment site, Cross Link attachment site) the number of pathogenic classifications is greater than or equal to the number of benign variants, but less than the number of variants of uncertain significance. Of these nine features, seven are single amino acid features, where, it appears, many changes may be less tolerated. The disruption of a disulfide bond by altering one of the cysteines involved, has the highest proportion of pathogenic variants. Of the reported 601 SNPs that affect disulfide bonds, 86% are pathogenic, 13% uncertain and 1% benign (see supplement Table S1), indicating that the resulting disruption of protein structure is very likely to be harmful. In comparison, variants that co-locate at single carbohydrate/glycosylation sites are tolerated best (less than 10% pathogenic assertions). The two types of features with the next highest proportion of pathogenic SNPs are Initiator methionine and Intramembrane region. Initiator methionine variants alter the initial methionine of a protein sequence, which is believed to result in the loss of protein translation. The Intramembrane region feature describes a sequence of amino acids entirely <u>within a membrane</u> but not crossing it.

**Figure 2.**
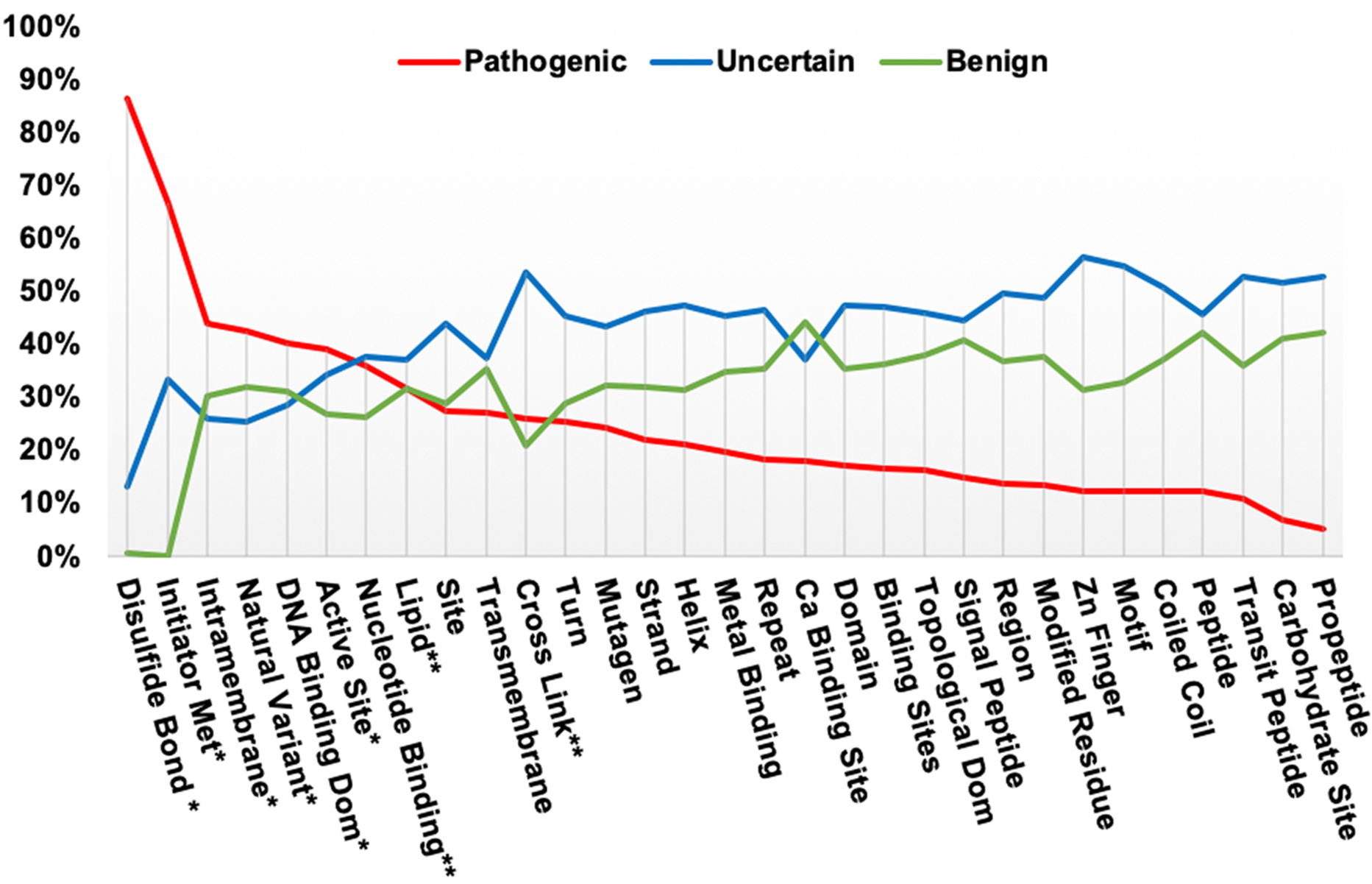
Percentage of ClinVar SNPs in each annotation category that exist in each feature type, underlying data table in supplemental methods. Features with “ * ” show greater pathogenic than benign or uncertain classifications. Features with “ ** ” have pathogenic classifications greater than or equal to benign, but less than uncertain.

### Comparison of ClinVar SNPs and UniProtKB Natural Variant Annotation

To survey genomic and protein annotations on variants we compared ClinVar SNPs with UniProtKB natural amino acid variations. In UniProt “Natural Variants” are polymorphisms including disease-associated mutations and RNA editing events. Currently 24,585 UniProt variants colocate on the genome with ClinVar SNPs (0-4 stars), which is 31.6% of all UniProtKB variants and 8% of all ClinVar SNP variants. Currently 53% of UniProtKB disease variants exist in ClinVar and 24.5% of ClinVar SNPs with pathogenic assertions are present as amino acid variants in UniProtKB. Table 2 shows a comparison between the UniProt variant classification of Disease, Unclassified and Polymorphism with ClinVar’s ACMG/AMP based assertions of ‘Pathogenic or Likely pathogenic’, ‘Uncertain Significance’ and ‘Benign or Likely benign’. The comparison in **Table 2** is a subset of all the co-located SNPs representing 35% of the total variants mapped as all ClinVar SNPs with 0 gold star evidence levels and some 1-star SNPs were excluded (see Methods). The table shows there is general agreement among similar annotations between the databases, with 86% of UniProtKB disease associated variants mapping to ‘pathogenic’ SNPs in ClinVar and with 10% falling into the middle ‘Uncertain Significance’ category. The remaining 4% fall mainly into the benign category. UniProt’s ‘Polymorphism’ category is closest in meaning to the ‘Benign’ categories in ClinVar; here, again, there is 85% agreement. For the remaining 15% of ‘Polymorphism’ variants 11% match the ‘Uncertain Significance’ category in ClinVar, 3% are classified as ‘pathogenic’ in ClinVar and 1% as ‘drug response’. UniProt’s ‘Unclassified’ category is closest in meaning to ClinVar’s ‘Uncertain Significance’; these are “grey” areas in each classification system and as such the agreement between the two databases is lowest: only 54% align and the rest are split between ‘pathogenic’ and ‘benign’ in ClinVar. The large number of variants annotated with ‘Uncertain Significance’ status is currently a general problem in the field (Hoffman-Andrews, 2017). In the ACMG/AMP framework, uncertain occupies a middle ground between benign and pathogenic. Often there is some evidence of a functional defect or harmful effect but it does not rise to clinical relevance or there is conflicting evidence.

**Table 2.** Mapping of Variants and annotation between ClinVar SNPs and UniProtKB amino acid variants that overlap in genome position and result in the same amino acid change. Only gold star rated ClinVar variants were included with evaluation criteria and no conflicts in assertions. Numbers in Bold face are comparisons discussed in the text. Numbers in parentheses are totals for each database.

|  | All ClinVar<br>SNPs<br>(249,784) | Pathogenic<br>SNPs<br>(27,819) | Uncertain<br>SNPS<br>(132,904) | Benign<br>SNPS<br>(94,541) | Other<br>SNPS<br>(11,489) |
| --- | --- | --- | --- | --- | --- |
| All UniProt Variants<br>(77,647) | **8609** | 3918 | 1291 | 3360 | 40 |
| Disease Variants<br>(30,220) | 4135 | **3562** | 412 | 159 | 2 |
| Unclassified<br>Variants (7,579) | 876 | 245 | **476** | 155 | 0 |
| Polymorphism<br>Variants (39,848) | 3598 | 111 | 403 | **3046** | 38 |

Though annotations in UniProt and ClinVar are in general agreement, there is still a significant level of disagreement between the databases, which is similar to that seen in recent analyses that compared variant pathogenicity interpretations by several laboratories within the ClinGen framework (Amendola et al., 2016; Gelb et al., 2018). These discrepancies may arise when variants have been assessed at different times, for different populations, and using different data types. Protein curators traditionally focus more on functional biochemical evidence to relate function to disease as well as evidence of genetic inheritance. In comparison, medical geneticists put more weight on genetic studies, variant frequencies, penetrance and, increasingly statistical models (InSiGHT; Plon et al., 2008) for variant classification

The most serious discrepancies seen here are the 270 variants (3%) that fall into the Pathogenic-Polymorphism and Benign-Disease categories (Table 2). UniProt curators are now able to investigate and correct or update the classification as required. At least 21 of these are clearly suspect as they have 3 stars in ClinVar indicating review by an ‘expert panel’. Twenty are variations in well-known oncogenes *MLH1, MSH2, MSH6, BRCA1,* and one in the *MYH7* gene is well known to be associated with cardiomyopathy. Inspection of the publications associated with the variants showed many more, publications associated with the ‘ClinVar submissions. Again, an example illustrates this situation. The variant P40692:p.Asp132His in the *MLH1* gene (uniprot.org/uniprot/P40692#VAR_022665) is classified as ‘disease’ in UniProt based on its association with colon cancer and experimental data in Nature Genetics (Lipkin et al., 2004). The equivalent SNP in ClinVar NM_000249.3(MLH1):c.394G>C (p.Asp132His) (https://www.ncbi.nlm.nih.gov/clinvar/variation/17096/) is annotated as ‘benign’ and reviewed by an ‘expert panel’ and cites the same 2004 study but also 14 additional more recent publications where the expert opinion has evolved.

Pharmacogenomic variants may also be classified differently by protein curators and medical geneticists. Our co-located data set in Table 2 has 52 variations with a ‘drug response’ annotation in ClinVar. Fourteen of these also have assertions of pathogenic, benign or uncertain, whereas 38 (0.6 %) have only ‘drug response’ and were all ‘reviewed by an expert panel’. For example, Q9BY32:p.Pro32Thr (uniprot.org/uniprot/Q9BY32#VAR 015576), a variant in the ITPA gene, is classified as a ‘disease’ variant in UniProt due to its association with heritable inosine triphosphatase deficiency. The UniProt annotation includes the notation: “It might have pharmacogenomic implications and be related to increased drug toxicity of purine analog drugs”. In ClinVar, the same variant, NM_033453.3(ITPA):c.94C>A (www.ncbi.nlm.nih.gov/clinvar/variation/14746/), is annotated by an expert panel at PharmaGKB (Whirl-Carrillo et al., 2012) as a “drug response” variant. The annotation cites literature about the variant’s effect on some antiviral drugs (Azakami et al., 2011; Chayama et al., 2011). Whilst the annotations in the two resources is different, both are correct based on the publications cited and each group’s area of interest.

### Comparison of Literature Citations

Positional mappings also allow comparison of literature cited as evidence for the annotated assertions. We compared all PMIDs cited as evidence for the co-located ClinVar and UniProt variants (the same set that was used for Table 2). Of these variants, 8,214 (95%) had a PMID in one or both databases, and 6,068 (70%) had one or more PMIDs in both databases. Of these, 4,001 (48%) shared one or more identical PMIDs. Not all variants cited a PMID: 7.6% have no PMIDs cited in ClinVar, 17.2% have none in UniProt, and 4.7% have no PMID listed in either database. In ClinVar, documentation, though encouraged, has not always been required for submission and documentation other than peer-reviewed publications is accepted. Also some ClinVar citations concern curation methods rather than the specific gene or variation. In UniProt, the missing PMIDs are an error. All the Natural Variants in the Swiss-Prot section were curated from literature cited in the entry. However, the link to the publications from the amino acid sequence is missing for some older and high throughput publications. Curation and data management practices changed years ago to solve this problem, but not all PMID links have been recovered.

Looking again at our biological example the *GLA* gene, 28 UniProt and ClinVar variants overlap and 27 of them agree on classification: 26 are classified as ‘Pathogenic/Likely pathogenic’ by ClinVar and Disease-associated by UniProtKB, one as “Unclassified” in UniProt and as “Uncertain significance, drug response” in ClinVar. Of the 28 common variations ClinVar has no citations for ten, and UniProtKB is missing citations for two. There is a total of 91 unique PMIDs: 81 from ClinVar and 10 from UniProtKB in the combined *GLA* set.

## Conclusion

Exome sequencing for clinical diagnosis is becoming more common and usually uncovers many non-synonymous SNP variations of unknown significance (VUS). Distinguishing which, if any, of these variants could be causal is difficult. Protein annotation can aid in variant curation by providing a functional explanation for a variant’s effect which is one of several important evidence categories used predicting the severity of variants (Nykamp et al., 2017; Richards et al., 2015). Accurate mapping between protein and genome annotation allows for more detailed analysis of the effects of variation on protein function. Here we have illustrated some of the types of comparative analyses that can be developed when this mapping information is available. The results in Figure 2 suggest that the location of a variant within some types functional features may be related to pathogenicity and might be useful in variant classification. For example, we observed that intramembrane regions, which do not include surface residues, show the highest number of variants in the pathogenic category for any multiple amino acid feature. In contrast, we did not observe more pathogenic variants in transmembrane regions, which cross the membrane but can contain some residues on the surface (Figure 2).

A recent comparison of 10,000 human genomes (Telenti et al., 2016) analysed a subset of 19,304 transmembrane regions from UniProtKB and observed less variation in the intramembrane region than outside the membrane boundary, suggesting a structural reason for amino acid conservation in those regions. Though interesting, the analyses are not directly comparable as Telenti et al. looked at variation in general inside and outside the membrane with a much larger set of variants and transmembrane regions. Here we looked at a smaller annotated ClinVar variant set, which only mapped to 6,754 (of a possible 43,734) transmembrane regions in 2,277 proteins. A more rigorous statistical analysis is needed that shows that a significant correlation of SNP pathogenicity within a particular protein functional feature. This analysis would need to account for potential confounding factors (e.g. feature size; overlapping features; and the quality, accuracy, and completeness of annotation). Then a feature specific correlation, could be a useful value to include in existing computation pipelines or new algorithms that evaluate and score variants.

The global comparison of variant classification between UniProtKB and ClinVar in Table 2 and the comparison of literature citations for variants between the two public databases were also informative. There is general agreement on classification of variants between genome and proteome even though priorities and terminology have been different. However, both comparisons illustrate that the classification of variations in enzymatic activities related to drugs needs better standardization. Many clinically relevant somatic variants found in tumors may need to be handled into a similar manner to ‘drug response’ variants, because they confer sensitivity or resistance to a treatment regime (Boca, 2018; Li et al., 2017; Madhavan, 2018; Ritter et al., 2016). Thus, the ‘pathogenic/benign’ terminology might not be appropriate for all cases.

The work described here provides the basis for a re-evaluation of UniProtKB annotation and the further standardization of this annotation with ClinVar and ClinGen. A detailed evaluation in which UniProt curators are performing a systematic re-curation of a randomly chosen set of variants from UniProt and ClinVar using the ACMG guidelines is being completed (M. Famiglietti et al., 2018). Recent efforts in the medical community to standardize the methods and levels of evidence required for the annotation of genetic variants (Amr et al., 2016; Manrai et al., 2016; O’Daniel et al., 2016; Richards et al., 2015; Walsh et al., 2016), along with increasing amounts of population data (Amr et al., 2016; Walsh et al., 2016), are leading to the widespread reevaluation of previous assertions of pathogenicity.

UniProtKB features have been mapped to the genome before, as the UCSC genome browser has provided selected UniProtKB/Swiss-Prot features for several years. The mappings described here contain additional annotation beyond that previously available and include isoform sequences from Swiss-Prot and sequences and features from the TrEMBL section of UniProtKB. The data files and track hubs will be updated with each release of UniProtKB, making any new annotation available immediately. The 34 features currently provided are not all of the positional annotations in UniProtKB, and we may add additional features in future releases. We plan to extend genome mapping to other model organisms. UniProt is working with the UCSC and Ensembl browser teams to improve the presentation of protein annotation on the respective browsers. In addition, some of the data provided here are available programmatically via a REST API (Nightingale et al., 2017). UniProt also collaborates with ClinVar to provide reciprocal links between variants that exist in both databases.

In summary, linking annotated data with assertions, publications and other evidence from UniProtKB, ClinVar or other datasets via co-location on the genome, as we demonstrate here, should help to better integrate protein and genomic analyses and improve interoperability between the genomic and proteomic communities to better determine the functional effects of genome variation on proteins. The location of a variant within functional features may correlate with pathogenicity and would be a useful attribute for use in variant prediction algorithms, including machine-learning approaches. We hope to investigate this and related topics in the future, and as a publicly funded resource, UniProt encourages others to further analyze our data as well.

## Data Access

The extended BED text files and binary BigBed files used for genome Track Hubs are available from the “Genome annotation tracks link in https://uniprot.org/downloads. Public Track hubs are available at the UCSC genome browser (Tyner et al., 2016) at (http://genome.ucsc.edu/cgi-bin/hgHubConnect?hubSearchTerms=uniprot) and the Ensembl genome browser (Aken et al., 2016; Hubbard et al., 2007) at (ensembl.org/Homo_sapiens/Info/Index#modal user data-Track Hub Registry Search) via a track hub registry search for “UniProt”. The Track Hub Registry (trackhubregistry.org/search?q=uniprot&type=genomics) provides links to view the links in either browser. Links from trackhubregistry.org that load the default UniProt tracks automatically are shown below. Additional tracks can be selected for display on each browser.

## UCSC Browser

genome.ucsc.edu/cgi-bin/hgHubConnect?db=hg38&hubUrl=ftp://ftp.uniprot.org/pub/databases/uniprot/current_release/knowledgebase/genome_annotation_tracks/UP000005640_9606_hub/hub.txt&hgHub_d_redir_ect=on&hgHubConnect.remakeTrackHub=on

## Ensembl Browser

www.ensembl.org/TrackHub?url=ftp://ftp.uniprot.org/pub/databases/uniprot/current_release/knowledgebase/genome_annotation_tracks/UP000005640_9606hub/hub.txt;species=Homosapie_ns;name=UniProtFeatures;registry=1

## Acknowledgments

Funding: NIH/NIGRI grants U41HG007822, U41HG007822-02S1 for UniProt and U01HG007437 for ClinGen. Additional UniProt funding sources can be found here: https://www.uniprot.org/help/about#funding

## Ethical Compliance

Patient clinical data have been obtained in a manner conforming with IRB and/or granting agency ethical guidelines.

