## Supplemental Methods and Figures for "UniProt Genomic Mapping for Deciphering Functional Effects of Missense Variants"

1. Innovation Center for Biomedical Informatics, Georgetown University Medical Center, 2115 Wisconsin Avenue NW, Suite 110, Washington, DC 20007, USA. 2. Center for Bioinformatics and Computational Biology, University of Delaware, Newark, DE 19711-5449, USA. 3. European Molecular Biology Laboratory, European Bioinformatics Institute (EMBL-EBI), Wellcome Genome Campus, Hinxton, Cambridge CB10 1SD, UK. 4. SIB Swiss Institute of Bioinformatics (SIB), Centre Medical Universitaire, 1 rue Michel Servet, 1211 Geneva 4, Switzerland; Protein Information Resource (PIR), Washington, DC and Newark, DE, USA; European Molecular Biology Laboratory, European Bioinformatics Institute (EMBL-EBI), Wellcome Genome Campus, Hinxton, Cambridge CB10 1SD, UK.

### Supplemental Methods

Mapping UniProtKB protein sequences to their genes and genomic coordinates is achieved with a four phase Ensembl import and mapping pipeline. The mapping is calculated for the UniProt human reference proteome with the GRC reference sequence provided by Ensembl. We intend to extend this mapping to yeast in the near future and other model organisms later. Here we provide additional details of the mapping calculation in Phase Three and additional information about the data fields available with the BED and BigBed files as explained in Phase Four. We also provide some additional detail of methods used to mapping positional features to ClinVar SNPs and how we compared UniProtKB variant annotations to ClinVar SNP annotation.

#### Phase Three: Converting UniProt Position Annotations to their Genomic Coordinates:

UniProt position annotations or “features” have either a single amino acid location or amino acid range within the UniProtKB canonical protein sequence. With the exon coordinates mapped to the protein peptide fragment, the genomic coordinates of a positional annotation are calculated by finding the amide (N) terminal exon and the carboxyl (C) terminal exon. The N-terminal or 5’ genomic coordinate ($UP_{Gcoord}$) of the positional annotation is calculated by 1. calculating the amino acid offset ($N_{aa}$) from the N-terminal protein peptide fragment start amino acid. 2. Taking the 5' genomic coordinate ($G_{coord}$) of the mapped exon and exon splice phasing (phase); the genomic coordinate is calculated as:

$$UP_{Gcoord} = G_{coord} + (N_{aa} * 3) + phase.$$

Likewise, the C-terminal, 3' genomic coordinate is calculated in the same way but with the 3' exon. For reverse strand mapped genes the above formula is modified to take into account the negative direction and phasing. Where a positional feature is spread out over multiple exons, the introns will be included in the mapping. This process is illustrated in **Figure 1**. If the positional feature is composed of a single amino acid, the three bases that denote that amino acid are given as the genomic coordinate. This is a limitation for the UniProt reviewed Natural Variants as UniProt does not independently define the specific allele change responsible for the missense, protein-altering variant. Therefore, UniProt is providing cross-references to dbSNP.

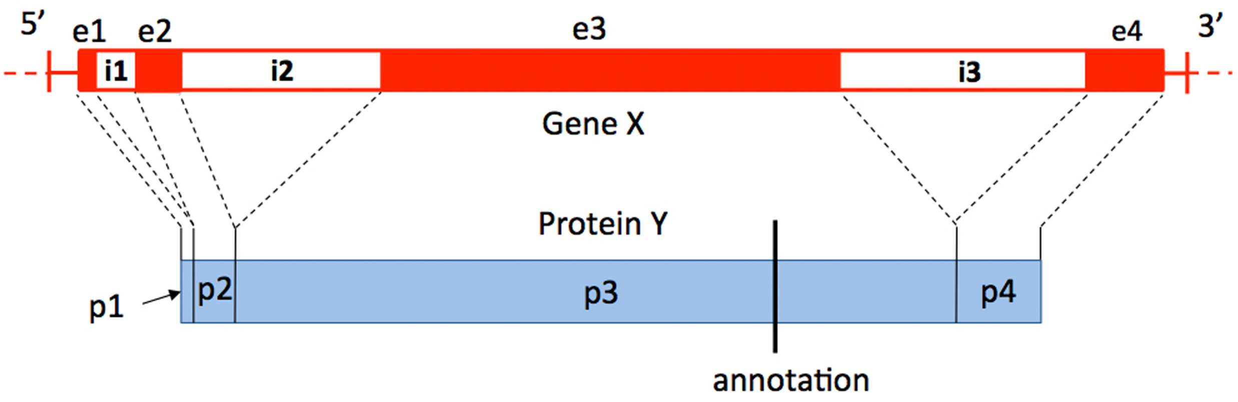


**Figure S1.** Converting UniProt Position Annotations to Genomic Coordinates. The genomic coordinates of a positional annotation are calculated by finding the N-terminal and the C-terminal of its exon. The genomic coordinate of the annotation in peptide p3 is calculated as:

$UP_{Gcoord} = G_{coord}$ of e3 start + $N_{aa}$ p3 peptide start to annotation * 3 + phase

Likewise, the C-terminal, 3' genomic coordinate is calculated in the same way but with the 3' exon.

**Phase four: UniProt BED and BigBed Files:** Converting protein functional information into its genomic equivalent requires standardized file formats. Genomic data is collated in files based upon a simple tab-delimited text format or the SAM (Sequence Alignment/Map) format (Li et al., 2009) The Browser Extensible Data (BED), a tab-delimited format, represents the best format type for converting UniProt annotations into genomic features for display in a genome browser. A BED file is interpreted as an individual horizontal feature 'track' when uploaded into a genome browser; this allows users to choose specific UniProtKB annotations most relevant to their analysis. The binary equivalent of the BED file is BigBed (W. J. Kent, Zweig, Barber, Hinrichs, & Karolchik, 2010). This file is more flexible in allowing for additional tab-delimited elements providing UniProt a greater opportunity to fully represent its protein annotations and one of the file formats used to make track hubs(Raney et al., 2014). A track hub is a web-accessible directory of files that can be displayed in track hub enabled genome browsers. Hubs are useful, as users only need the hub URL to load all the data into the genome browser. Moreover, a public registry for track hubs is now available (https://trackhubregistry.org/) allowing users to search for track hubs directly through the genome browser rather than searching for hubs at the institute or bioinformatics resource that generated the data.

Using the protein and genomic coordinates with additional feature specific annotations from UniProtKB, BED (UCSC, 2016a) and BED detail (UCSC, 2016b) formatted files were produced for the UniProtKB human reference proteome by converting the genomic mappings to the zero-based genomic coordinates used by the BED format. A region of DNA (or block), its size and offset from the start of DNA being annotated is calculated for a protein annotation or sequence. If a protein annotation or sequence is defined by a range (e.g. chain, domain, region) or is composed of more than one amino acid or sub-region (e.g. exon) one block element is defined in a comma separated list for each sub-region (e.g. each exon). This means that a protein annotation could be represented as a single block or more than one block depending upon its composition. Blocks and block sizes for the UniProt proteome sequences define the specific exons and the sizes of those exons that are translated into the protein sequence. A standard BED file is generated for the proteome sequences and BED detail files are generated for each positional annotation type listed in Table S1, except Natural Variant. In both file formats the UniProtKB accession for the sequence or individual annotation is provided in the BED name column (column 4) to provide a convenient link to the original UniProt entry. In the BED detail files, the last two columns are used for UniProtKB annotation identifiers and description, when available. The description is composed of the protein position or range of the annotation within the protein, any functional description and any literature evidence associated to the annotation. Variant BED file differs in defining the protein HGVS for the variant instead of the annotation position or range.

UniProt BigBed files differ from the BED detail files with the addition of extra columns that separate the final description column of the BED detail file. For all UniProt positional annotations columns 13 and 14 become an additional entry identifier field and annotation type field, respectively. Then for all annotation types, except variant, four additional columns are used to define an annotation identifier, annotation position or range, description and literature evidence. Variant BigBed files differ from the other annotation type BigBed files by changing column 17, description, to a disease description and column 18, literature evidence, to a protein HGVS representation of the variant. Then a further three columns are defined; one: any variant cross-references, eg the variant is also reported in ClinVar, two: any general description about the variant and finally, literature evidence for the variant. BigBed files (W. J. Kent et al., 2010) are produced from the text BED detail files using the UCSC bedToBigBed converter program (http://hgdownload.cse.ucsc.edu/admin/exe/). bedToBigBed requires additional information defining the column structure of the BED detail file as an autoSQL (J. Kent & Brumbaugh, 2002) (http://hgwdev.cse.ucsc.edu/~kent/exe/doc/autoSql.doc) and the chromosome names and sizes for the genome assembly where UniProt use the chromosome names and sizes available from Ensembl's latest assembly release. Binary BigBed files are generated for each type sequence annotation and the UniProtKB human proteome protein sequence set.

**Mapping ClinVar SNPs to protein features and variants**: Data for comparing ClinVar SNPs to UniProt features comes from the ClinVar variant_summary file from NCBI (ftp.ncbi.nlm.nih.gov/pub/clinvar/tab_delimited/variant_summary.txt.gz), the UniProtKB feature specific BED files (ftp.uniprot.org/pub/databases/uniprot/current_release/knowledgebase/genome_annotation_tracks/UP000005640_9606_beds/) and the human variation file on the UniProt FTP site: (ftp.uniprot.org/pub/databases/uniprot/current_release/knowledgebase/variants/humsavar.txt). 1) For each feature in UniProtKB, we check the genomic position against the position for each record in ClinVar. If the genome positions of the protein feature overlap the chromosome and genomic coordinate of the SNP we establish a mapping. Information about the SNP and the feature, including the amino acid change are attached to the mapping file. 2) For each result in 1, we check the SNP position against the exon boundary for the Protein. A flag is added if a SNP coordinate is within the exon boundary. Variants outside of exons were excluded from further analysis. 3) The disulfide bond feature is two distinct Cys residues who form a bond causing a peptide loop between them but for historical purposes is annotated in UniProt as a range between the Cysteines. This required some extra processing of this feature to remove all but the first and last Amino Acids from the comparison with ClinVar. 4) For each UniProt variant in 2 and 3, we check that the ClinVar provided and UniProt accession numbers refer to the same protein and also that the specific amino acid changes reported in UniProt and ClinVar is the same. 5) To compare pubmed IDs (PMIDs) cited as evidence in ClinVar we used the file (ftp.ncbi.nlm.nih.gov/pub/clinvar/tab_delimited/var_citations.txt) from NCBI and the UniProtKB/Swiss-Prot file (ftp.uniprot.org/pub/databases/uniprot/current_release/knowledgebase/complete/uniprot_sprot.dat.gz) from Nov 2018. For each collocated variant we used the ClinVar Allele ID and UniProtKB Variant ID to extract the relevant PMIDs for each variant and constructed a table of all the PMIDs and counts of the PMIDs for the collated Allele IDs and Variant IDs.

**Comparison of UniProt and ClinVar Variant Annotation:** UniProt curators classify variants into three categories: 1) Disease - variants reported to be implicated in disease; 2) Polymorphism - variants not reported to be implicated in disease; 3) Unclassified - variants with uncertain implication in disease as evidence for or against a pathogenic role is limited, or reports are conflicting. ClinVar does not annotated variants directly but accepts submitters assertions of clinical significance with their criteria and classifies them into 0-4 gold star groups based on levels of evidence. The predominant assertions in ClinVar and the ones we used for comparison are those recommended by the ACMG/AMP guidelines (Richards et al., 2015) Benign, Likely benign, Uncertain significance, Likely pathogenic and Pathogenic. In addition, there are a small number of disease related assertions in ClinVar labled as ‘risk factor’ and ‘drug response’. For our comparison we only used variants with 1-4 stars but removed all 1-star variants with conflicting interpretations and those with no associated phenotype, for example they contained only MedGen codes CN169374 (not specified) and/or CN517202 (not provided). We compared ClinVar assertions to UnProt classifications as follows: all 'pathogenic' assertions (pathogenic and likely pathogenic) to 'Disease' in UniProt; 'Uncertain significance' with 'Unclassified'; and, all 'benign' (benign and likely benign) assertions to 'Polymorphism'. Anything else in ClinVar was grouped as 'other' in the results. All the 'others' that aligned with UniProt annotations in this comparison were ClinVar 'drug response' assertions.

**Supplemental Results.**

**Biological Example 2:** **Figure S2** shows a portion of the *APP* gene that encodes the amyloid beta A4 protein (UniProtKB P05067) on the Ensembl genome browser. Amyloid beta A4 protein is a cell surface receptor with multiple functions related to neurite growth. Peptides derived from cleavage of the mature protein are found in amyloid plaques in brain tissue from patients with Alzheimer disease (Ancolio et al., 1999; Chartier-Harlin et al., 1991; Cras et al., 1998; Denman, Rosenzcwaig, & Miller, 1993; Eckman et al., 1997; Goate et al., 1991; Hendriks et al., 1992; Kwok et al., 2000; Liepnieks, Ghetti, Farlow, Roses, & Benson, 1993; Mullan et al., 1992; Murrell, Farlow, Ghetti, & Benson, 1991; Nilsberth et al., 2001). Cleavage into multiple chains and peptides occurs at eight cleavage sites by $\alpha$-, $\beta$-, $\gamma$- and $\theta$- secretase and various caspase enzymes. Non-amyloid forming proteolysis occurs with $\alpha$ and $\gamma$ secretases resulting in alpha species sAPP$\beta$ and a C-terminal fragment ($\alpha$CTF). **Figure 2** shows two $\gamma$-secretase cleavage sites between positions 711-712 and 713-714 on the protein responsible for producing the N terminals of $\beta$-APP42 and $\beta$-APP40 peptides. The $\beta$-APP40 form aggregates to generate amyloid beta filaments that are the major component of the toxic amyloid plaques found in the brains of Alzheimer disease (AD) sufferers (Selkoe, 1998). Disease-associated variants annotated in UniProtKB and ClinVar align with the cleavage sites, suggesting that alteration of cleavage can affect disease progression. The associated annotation shows that variant P05067:p.Thr714Ile (uniprot.org/uniprot/P05067#VAR_014218) was found in a family showing autosomal dominant inheritance of early-onset Alzheimer disease. The variation resulted in an ~11-fold increase in the $\beta$-APP42/$\beta$-APP40 ratio *in* *vitro* as measured by multiple methods and coincided with deposition of nonfibrillar pre-amyloid plaques composed primarily of N-truncated β-APP42 in the brain (Kumar-Singh et al., 2000).

**Supplemental Figures and Tables.**

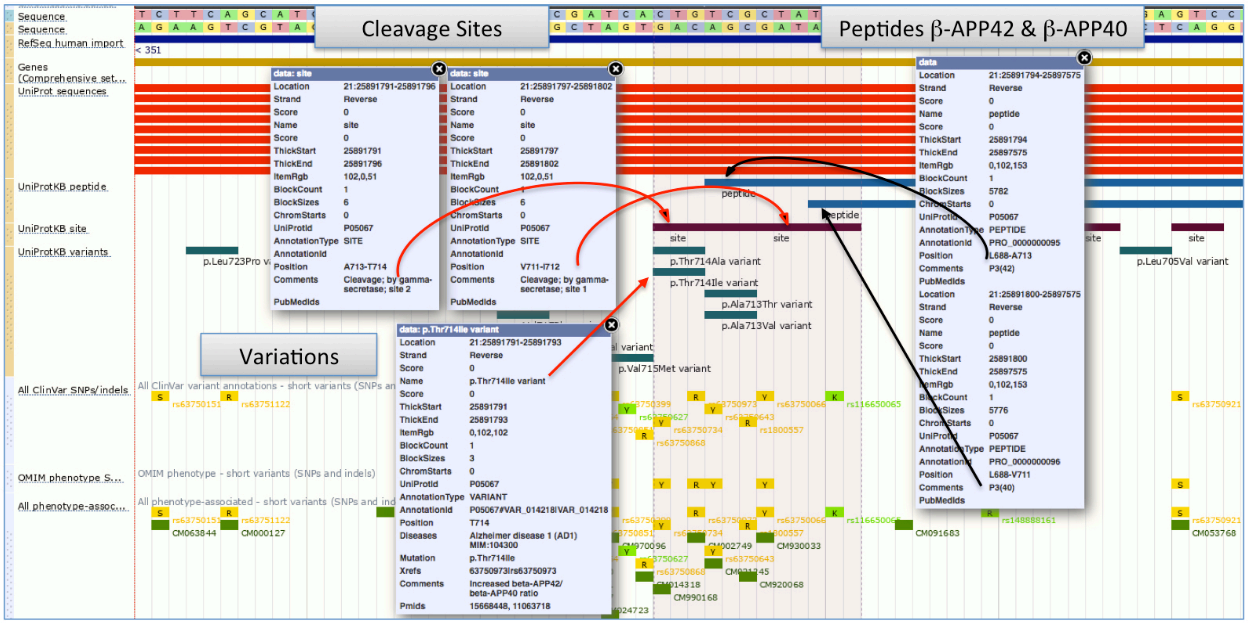


**Figure S1.** The *APP* gene (amyloid beta A4 protein, UniProt Acc. P05067) associated with Alzheimer disease shown on the Ensembl browser with selected UniProt Genome Tracks. Shows how pathogenic variations overlap protein cleavage sites contributing to the aberrant ratios of the β-APP42 and β -APP40 peptides observed in some Alzheimer cases.

**Supplemental Table S1**

| UniProt Feature Type | ClinVar SNPs in feature | Pathogenic SNPs in feature | Uncertain SNPs in feature | Benign SNPs in feature | % Pathogenic | % Uncertain | % Benign |
|---|---|---|---|---|---|---|---|
| Disulfide Bond* | 601 | 519 | 78 | 4 | 86% | 13% | 1% |
| Initiator Met* | 36 | 24 | 12 | 0 | 67% | 33% | 0% |
| Intramembrane* | 208 | 91 | 54 | 63 | 44% | 26% | 30% |
| Natural Variant* | 26,890 | 11,416 | 6,818 | 8,596 | 42% | 25% | 32% |
| DNA Binding Dom* | 1,343 | 541 | 384 | 418 | 40% | 29% | 31% |
| Active Site* | 41 | 16 | 14 | 11 | 39% | 34% | 27% |
| Nucleotide Binding* | 625 | 224 | 235 | 164 | 36% | 38% | 26% |
| Lipid* | 184 | 58 | 68 | 58 | 32% | 37% | 32% |
| Site | 523 | 143 | 229 | 150 | 27% | 44% | 29% |
| Transmembrane | 11,318 | 3,077 | 4,230 | 3,994 | 27% | 37% | 35% |
| Cross Link* | 58 | 15 | 31 | 12 | 26% | 53% | 21% |
| Turn | 1,356 | 343 | 614 | 392 | 25% | 45% | 29% |
| Mutagen | 814 | 198 | 352 | 263 | 24% | 43% | 32% |
| Strand | 8,508 | 1,858 | 3,931 | 2,710 | 22% | 46% | 32% |
| Helix | 11,605 | 2,440 | 5,504 | 3,645 | 21% | 47% | 31% |
| Metal Binding | 3,144 | 619 | 1,427 | 1,096 | 20% | 45% | 35% |
| Repeat | 14,618 | 2,665 | 6,786 | 5,160 | 18% | 46% | 35% |
| Ca Binding Site | 277 | 50 | 103 | 122 | 18% | 37% | 44% |
| Domain | 131,503 | 22,572 | 62,376 | 46,420 | 17% | 47% | 35% |
| Binding Sites | 10,943 | 1,812 | 5,144 | 3,975 | 17% | 47% | 36% |
| Topological Dom | 25,737 | 4,147 | 11,775 | 9,786 | 16% | 46% | 38% |
| Signal Peptide | 3,114 | 458 | 1,382 | 1,273 | 15% | 44% | 41% |
| Region | 36,621 | 5,023 | 18,154 | 13,426 | 14% | 50% | 37% |
| Modified Residue | 742 | 100 | 361 | 280 | 13% | 49% | 38% |
| Zn Finger | 2,518 | 312 | 1,418 | 788 | 12% | 56% | 31% |
| Motif | 799 | 98 | 438 | 263 | 12% | 55% | 33% |
| Coiled Coil | 14,447 | 1,769 | 7,310 | 5,362 | 12% | 51% | 37% |
| Peptide | 3,931 | 478 | 1,794 | 1,658 | 12% | 46% | 42% |
| Transit Peptide | 459 | 50 | 242 | 165 | 11% | 53% | 36% |
| Carbohydrate Site | 163 | 11 | 84 | 67 | 7% | 52% | 41% |
| Propeptide | 784 | 40 | 414 | 330 | 5% | 53% | 42% |

Table S1. ClinVar SNPs that overlap UniProt Features. Only 1-4 gold star rated ClinVar variants were included that had evaluation criteria and no conflicts in pathogenicity assertions. Comparison uses UniProt and ClinVar January 2018 releases. Percentages of some features may not sum to 100%. This table was used to create Figure 2 in the main manuscript.
